# The rhythms of predictive coding: pre-stimulus phase modulates the influence of shape perception on luminance judgments

**DOI:** 10.1101/061309

**Authors:** Biao Han, Rufin VanRullen

## Abstract

Predictive coding is an influential model emphasizing interactions between feedforward and feedback signals. Here, we investigated its temporal dynamics. Two gray disks with different versions of the same stimulus, one enabling predictive feedback (a 3D-shape) and one impeding it (random-lines), were simultaneously presented on the left and right of fixation. Human subjects judged the luminance of the two disks while EEG was recorded. Independently of the spatial response (left/right), we found that the choice of 3D-shape or random-lines as the brighter disk (our measure of post-stimulus predictive coding efficiency on each trial) fluctuated along with the pre-stimulus phase of two spontaneous oscillations: a ~5Hz oscillation in contralateral frontal electrodes and a ~16Hz oscillation in contralateral occipital electrodes. This pattern of results demonstrates that predictive coding is a rhythmic process, and suggests that it could take advantage of faster oscillations in low-level areas and slower oscillations in high-level areas.

## Introduction

The outside world provides us only the light, but our visual system is capable of extracting the basic features in low-level areas and understanding them as meaningful concepts in high-level areas. Predictive coding theory suggests that the brain employs an efficient coding strategy to achieve this by generating predictions in higher-level areas and comparing them with the incoming sensory signals in the lower-level areas (Friston, 2005; Rao and Ballard, 1999). Previous neuroimaging evidence revealed the existence of such two-way communication (Alink et al., 2010; Egner et al., 2010; Harrison et al., 2007; Murray et al., 2002; Summerfield et al., 2008). However, the underlying mechanisms in this dynamical process, especially in the temporal domain, remain unknown.

It has been proposed that the feed-forward and feedback in predictive coding take advantage of oscillations for information processing (Bastos et al., 2015; Fontolan et al., 2014). On the one hand, recent neurophysiological evidence on laminar-specific oscillations and the functional roles of different layers suggested a faster oscillation for the feed-forward pathway and a slower oscillation for the feedback pathway (Bastos et al., 2012; Buffalo et al., 2011; Fontolan et al., 2014; Maier et al., 2010; van Kerkoerle et al., 2014). On the other hand, recent studies showed a link between behavioral performance and cortical oscillations in perception (Busch and VanRullen, 2010; Dugué et al., 2011) and reaction time (Drewes and VanRullen, 2011; Song et al., 2014). Since neural oscillations can reflect the cyclic fluctuations of excitability in a network, investigation of the relationship between trial-to-trial variability and the phase of ongoing oscillations could link specific oscillations to cognitive functions (e.g. attention).

Here, we used this approach to investigate the specific influence of ongoing oscillations on predictive coding, by measuring its effect on perception for different pre-stimulus oscillatory phases. In a typical predictive coding experiment, two conditions must be created: one with strong prediction signals, one without. Since the predictions are sent via feedback signals to lower areas, they will affect the lower-level activity and thus presumably also affect perception. Here, we chose one of the first paradigms in predictive coding to generate different amounts of predictive feedback (Murray et al., 2002): shape perception.

Specifically, 3D-shape outlines and random-lines versions of the same stimuli, similar to the stimuli used in a previous influential study (Murray et al., 2002), were used in this experiment. It has been shown that 3D-shape outlines can be easily recognized and thus produce more predictive feedback than the random-lines versions (Murray et al., 2002). To measure the effect of different amounts of predictive feedback, we asked the subjects to judge the luminance (report the side of the brighter disk) of two gray disks simultaneously displayed on the left and right of fixation on a black background, one containing the 3D-shape outlines and the other containing the random-lines version. The luminance of the disks was adjusted to achieve about 50% choice rate for 3D-shape/random-lines disk (this was achieved by slightly increasing the luminance of the random-lines disk, as demonstrated in one of our previous studies (Han and VanRullen, 2016, 2014)). We recorded EEG signals and analyzed the relationship between pre-stimulus oscillation phase and the post-stimulus judgment. If predictive feedback mechanisms during the post-stimulus stage involved one or more periodic processes, then the oscillatory state of the system just before stimulus onset (as reflected in pre-stimulus oscillatory phase) should have an influence on the outcome of predictive feedback, and thus on the perceptual judgment. We found that, independent from the spatial choice (left/right side), the phase of 5Hz contralateral frontal and 16Hz contralateral occipital pre-stimulus oscillations modulated the subject's choice of a brighter 3D-shape disk (more effective predictive feedback) or a brighter random-lines disk (less effective predictive feedback). Since higher hierarchical level areas are assumed to send predictive feedback and lower hierarchical level areas to send prediction error, our results imply that in the post-stimulus stage the brain sends predictive feedback periodically at a preferred phase of a theta frequency oscillation in the frontal region, and sends prediction errors periodically at a preferred phase of a beta frequency oscillation in the occipital region.

## Materials and Methods

### Subjects

Fifteen volunteers participated in the experiment. One participant was excluded from the analysis due to the poor behavioral performance in catch trials (<60% trials were correctly reported, with a chance level of 50%, see below). Fourteen participants remained in the sample (8 female, mean age 28.01 ± 4.81 years, four left-handed, four with left eye dominance). All subjects had normal or corrected to normal vision. The study was approved by the local ethics committee “Sud-Ouest et Outre-Mer I” and followed the Code of Ethics of the World Medical Association (Declaration of Helsinki). All subjects provided signed informed consent before starting the experiments.

### Apparatus

Stimuli were presented at 57 cm distance using a desktop computer (2.09 GHz Intel processor, Windows XP) with a cathode ray monitor (resolution: 800×600 pixels; refresh rate: 140 Hz, Gamma corrected luminance function). Stimuli were designed and presented via the Psychophysics Toolbox (Brainard, 1997) running in MATLAB (MathWorks).

### Stimuli and tasks

Stimuli consisted of a central white fixation point (diameter: 0.2 degrees of visual angle) and two circular gray disks (diameter: 4 degrees each) presented randomly to the left and right of fixation (3 degrees eccentricity). One 3D-shape stimulus was in the center of one disk (3D-shape disk) and one random-lines version of the same stimulus in the other (random-lines disk). The 3D-shape and random-lines stimulus pair was randomly chosen from twenty pairs of stimuli generated beforehand using a method similar to Murray et al. (2002): 3D-shapes were generated by randomly selecting 4-6 vertices, connecting the vertices and adding small extensions to render perceived depth; random-lines stimuli were created by breaking the 3D-shape at its intersections and randomly shifting the lines within the display (Murray et al., 2002). The diameter of both 3D-shape and random-lines stimuli was 3 degrees. The stimulus outlines were black.

Before stimulus onset, there was a blank screen with only the fixation point that lasted from 1000 to 1500ms (random uniform distribution). Then the two disks and the fixation point appeared for 150ms. After that, a question mark appeared in the center of the screen. There were two kinds of randomly mixed experimental trials: the main experimental trials and the catch trials. In main experimental trials, the luminance of the disks was adjusted (i.e. the random-line disks were set 1.45% brighter than the 3D-shape disks) based on a previous study (Han and VanRullen, 2016, 2014) to obtain an average 50% selection rate of 3D-shape/random-lines disks. In catch trials, one of the disks had its luminance value changed up or down by 20% compared to the luminance used in the main experimental trials, while the other disk kept the same luminance as in the main experimental trials. Subjects were presented with 4 or 8 blocks of 200 trials with 85% main experimental trials and 15% catch trials (the first 6 of the 14 subjects performed only 4 blocks of the present experiment, together with 4 blocks of another experiment that was eventually canceled and whose data were not analyzed). Subjects were instructed to fixate the fixation point all the time, judge the luminance of the disks and respond using the arrow keys (left arrow to indicate that left disk is brighter, right arrow for right disk brighter) on a standard 105 key keyboard when the question mark appeared. There was no feedback after the response.

### EEG data acquisition and analysis

EEG was recorded at 1024 Hz using a Biosemi system (64 active electrodes). Horizontal and vertical electro-oculograms (EOG) were recorded by three additional electrodes around the subjects' eyes. For data pre-processing, the EEG and EOG data were downsampled offline to 256 Hz, re-referenced to average reference and epoched around the stimulus onset in each trial for data analysis *via* the MATLAB (MathWorks) and EEGLAB toolbox (Delorme and Makeig, 2004). Individual electrode data were visually inspected, and channel data containing artifacts were interpolated by the mean of adjacent electrodes (three subjects had one electrode containing artifacts, one subject had two; the positions of the interpolated electrodes were different across subjects).

As the post-stimulus spatial choice was lateralized on each trial to the left or right side, the pre-stimulus oscillatory correlates of the post-stimulus luminance judgment may not only reflect the oscillation's influence on shape perception and predictive coding, but also its influence on spatial choice (i.e., pre-stimulus oscillations may bias the left/right spatial choice independently of the 3D-shape/random-lines content inside of the disk). To avoid any contribution from the spatial choice, we first divided the trials for each subject into two trial datasets based on the post-stimulus spatial choice, and performed the time-frequency analysis (described below) within each dataset. We reasoned that this analysis would lead to shape perception correlates not on a given fixed set of electrodes, but rather on different electrode groups depending on the side of choice (i.e., electrodes “contralateral” or “ipsilateral” to the spatial choice). Therefore, we arbitrarily chose to permute the electrode locations for the dataset corresponding to a right-side choice: we replaced the left-hemisphere electrodes by the symmetric ones from the right and vice versa (midline electrodes were unaffected). With this new electrode assignment, left-hemisphere electrodes would thus always correspond to those ipsilateral to the spatial choice, and right-hemisphere electrodes to contralateral ones.

For the time-frequency analysis, time-frequency transformations were first generated over all channels using EEGLAB with a function akin to a wavelet transform, starting with 3 cycles at 2Hz and increasing to 5 cycles at 50 Hz in the multiple-cycle analysis, and with 1 cycle from 2Hz to 50 Hz in the one-cycle analysis. This yields a complex representation of the amplitude, *A*, and the phase, φ, for trial *j* at time *t* and frequency *f*:

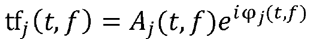

The phase of this representation can be extracted by normalizing the complex vector to the unit length:

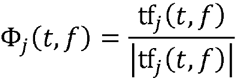

Inter-trial phase coherence (ITPC) measures the phase consistency across trials. We calculated the ITPC using the method described previously (Lachaux et al., 1999):

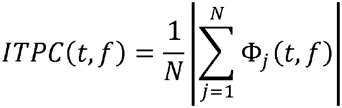

where N is the number of trials in one group of trials.

Here, we wanted to evaluate the relation between the pre-stimulus oscillatory phase and the influence of shape perception on luminance judgment (our measure of the efficiency of predictive coding). Would a particular pre-stimulus phase occur more frequently for trials with post-stimulus 3D-shape disk choice, and the opposite phase for trials with post-stimulus random-lines disk choice? In the pre-stimulus period, because intertrial intervals are randomized and unpredictable, the phase of the spontaneous EEG signal at a given prestimulus time should follow a uniform distribution over all trials. However, if there is a systematic relation between EEG phase and behavioral outcome, higher-than-chance inter-trial phase coherence should be observed in each of the trial subgroups (If EEG phases of two subgroups corresponding to different behavioral outcomes tend to oppose each other, the phases of one subgroup will concentrate around one phase angle and the phases of the other subgroup will concentrate around the other phase angle. Thus, high ITPC values will be observed for both subgroups. *Vice versa*, if EEG phases of two subgroups do not oppose each other, the phases of the two subgroups will distribute uniformly across different phase angles. Thus, low ITPC values will be observed. Therefore, the ITPC values are reliable indicators of phase opposition.). In that case, the product or the sum of the two subgroup inter-trial phase coherence could summarize, in a single variable, the strength of the phase-behavior relation (Busch et al., 2009; VanRullen et al., 2011). Here we thus introduce a new measure of the phase-behavior relation: *Phase Opposition Product (POP)*. This measure is calculated using the product of the inter-trial phase coherence of different trial subgroups:

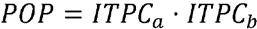

The measure of POP will be maximal when two subgroups show strong inter-trial phase coherence, in other words, strong phase opposition between two subgroups. To accurately assess the significance of the phase-behavior relation without any assumption about the probability distribution of the POP values, we performed a nonparametric permutation test: We first computed the POP values for each point in the time-frequency plane from −650 to 150ms, from 2 to 50 Hz for each electrode, dataset, and subject and then averaged across all datasets and all subjects. Surrogate POP values were obtained by randomly assigning the trials to one or the other condition for each subject (keeping the number of trials in each condition constant) and recalculating the grand-average POP values. We computed the P value by simply counting the number of surrogate POP values that were more extreme than the observed value. Here, we used 80,000,000 surrogates and thus assigned the P value of 1.25 × 10^-8^ to the points without any more extreme POP values in the surrogates. The P values were corrected for multiple comparisons over time points, frequencies and electrodes using the FDR method (FDR α=0.05, corresponding to a P value threshold of 9.53×10^-6^). To show the overall POP in the time-frequency domain, we computed a z-score by combining the observed POPs across all datasets, subjects, and electrodes and comparing the value with the mean and SD of a null-hypothesis distribution with 10,000 surrogate POP values (generated using the procedure described before, and also combined across all electrodes, subjects, and datasets).

## Results

Human observers judged the luminance of two disks that were presented for 150ms on the left and right of a central fixation point. The disks contained different versions of the same stimulus, one with a 3D-shape enabling predictive feedback, and the other with a random-lines version of the same shape which impeded predictive feedback (Figure 1). Before the stimulus onset there was a random period of time (1000-1500ms) with only the fixation point on the screen. After the stimulus offset a question mark appeared at the center and the subjects were instructed to report the side with the brighter disk. In the main experimental trials, the luminance of the disks was adjusted, such that observers reported the 3D-shape disk as brighter in half of the trials. 15% of trials were catch trials: extreme luminance values were assigned to one disk to monitor the subject's ability to judge the luminance difference.

**Figure 1.**
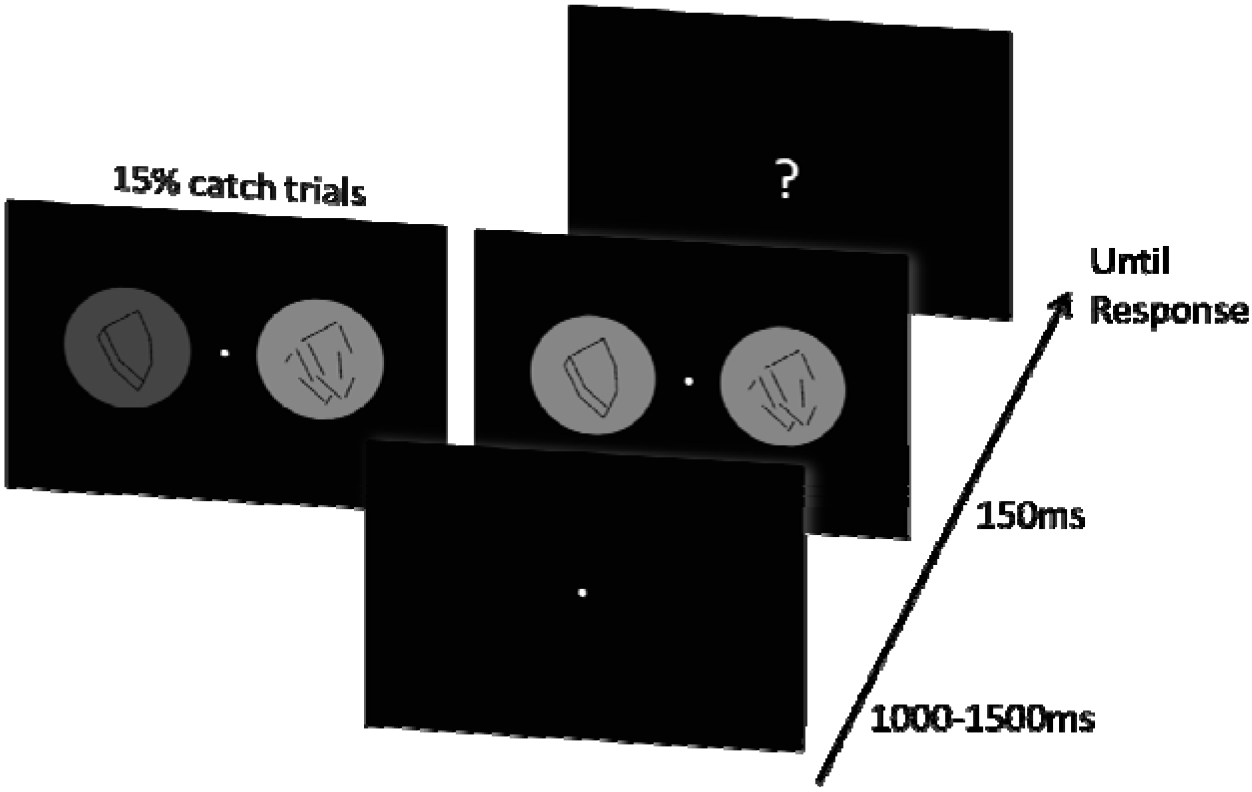
Illustration of the experimental paradigm. In each run of trials, a blank screen with only a central fixation point was presented for 1000 to 1500ms randomly. Then, two circular gray disks, one with a 3D-shape stimulus in the center and the other with random-lines, were presented randomly on either side (left or right) of the fixation point for 150ms. Subsequently, a question mark appeared in the center of the screen. Subjects were instructed to fixate the fixation point all the time, and report the side of the brighter disk with the corresponding arrow key after the question mark appeared. In the main experimental trials, luminance values of the disks were adjusted to obtain a 50% selection probability of 3D-shape/random-lines disk. In addition, there were 15% catch trials intermixed with the main experimental trials to monitor the subjects' ability to judge the luminance difference throughout the experiment. In these catch trials, one disk was 20% brighter/darker than in the main experimental trials.

**Behavioral Results.** On average, subjects judged the 3D-shape disk as brighter in half of the trials (49.24% ± 1.57%, mean ± standard error of the mean, SEM) in the main experimental condition, as expected. The luminance judgment correct rate in the catch trials (subjects judged the disk with higher luminance value as brighter or judged the disk with lower luminance value as darker) was high (93.98% ± 1.83%, mean ± SEM), indicating that subjects were adequately engaged in the luminance judgment task.

**Electrophysiological Results.** We focused on the relationship between pre-stimulus oscillatory phase and the trial-by-trial variations in the efficiency of post-stimulus predictive coding. EEG was recorded during the experiment. We expected the relation between oscillatory phase and behavior to be most visible in the pre-stimulus time window, where phase information reflects spontaneous fluctuations in neuronal excitability (Bishop, 1932; Busch et al., 2009; Buzsáki and Draguhn, 2004; Fries et al., 2007; Jensen et al., 2012). In contrast, post-stimulus phase information is driven to a large extent by stimulus-locked activity (e.g. evoked potentials) and is thus further removed from spontaneous activity. We used classical stimuli (Murray et al., 2002) for inducing different amounts of predictive feedback on the left and right of the screen: 3D-shape and random-lines versions of the same stimuli (Figure 1; the 3D-shape version enabling predictive feedback, the random-lines simultaneously impeding it). To measure the effective amount of post-stimulus predictive feedback on each trial, we probed the perceived luminance of the disks under the stimuli. We have previously demonstrated that the net effect of predictive feedback on these stimuli is a relative increase of perceived luminance for the disk containing the 3D-shape (Han and VanRullen, 2016, 2014). Here, this net effect was compensated on each trial by slightly lowering the luminance of that disk so that the average likelihood of perceiving either disk brighter was about 50% (see Methods); therefore, residual fluctuations of luminance perception on every trial can be thought to arise from trial-by-trial fluctuations in the efficiency of post-stimulus predictive coding (the 3D-shape disk may still be perceived brighter on trials where predictive coding was more efficient than average, and darker on trials where it was less efficient than average). Of course, spatial bias and/or trial-by-trial fluctuations in the direction of spatial attention may well also contribute to the choice of which disk appears brighter on a given trial. Thus, for each subject we divided all trials into two datasets based on their spatial choice (left-side choice vs. right-side choice), and we only investigated the relation between pre-stimulus EEG phase and 3D-shape/random-lines choice within each dataset. As the correlates of choosing the prediction-consistent stimulus (3D-shape) were expected to be strongest on electrodes contralateral to that stimulus, which would map onto opposite hemispheres for the two datasets, before plotting any scalp topographies we permuted the electrode positions (symmetrically across the midline axis) of all right-side choice trials. This procedure resulted in a mapping of ipsilateral effects to the spatial choice onto left electrodes, and contralateral effects onto right electrodes.

We estimated the relation between EEG phase and predictive coding via the phase opposition product (POP, see Methods). This measure should be maximal when 3D-shape choice trials (prediction-consistent) and random-lines choice trials (prediction-inconsistent) tend to have opposite phase values. For each subject, dataset, electrode, time point and oscillatory frequency, we obtained surrogate POP values (80,000,000 surrogates) by randomly permuting the trial outcomes, keeping the number of trials constant. Both real and surrogate POP values were averaged across datasets and subjects. The significance was determined as the proportion of surrogate POP values that were more extreme than the observed value. P-values were corrected for multiple comparisons across time points, frequencies and electrodes (100×30×64) using the FDR method (FDR α=0.05, corresponding to a P value threshold of 9.53 × 10^-6^). To show the overall POP in the time-frequency domain, a z-score was computed by comparing the real POP values (combined across all subjects, datasets, and electrodes) to the mean and standard deviation of a null-hypothesis distribution with 10,000 surrogate POP values (generated using the same procedure described before, and also combined across all subjects, datasets, and electrodes). This analysis revealed a significant phase-behavior relation between the post-stimulus 3D-shape/random-lines choice and two pre-stimulus oscillations (Figure 2. A): one theta-frequency oscillation (~3.1 Hz to 7.6 Hz) in the time window from −545 ms to −268 ms, and one beta-frequency oscillation (~13.2 Hz to 25.7 Hz) in the time window from −107 ms to −25 ms. Green outlines mark the significant time-frequency regions (at least one significant electrode) after FDR correction.

Scalp topographies of the z-score show that the two oscillatory effects involve distinct electrode groups and presumably distinct brain regions (Figure 2. B and C): the theta-frequency effect is maximal over frontal regions and the beta-frequency effect over occipital regions. In both cases, these effects are contralateral to the side that subjects chose as “brighter” (i.e., the right side of the topographies, due to our electrode permutation procedure). Electrodes with at least one significant time-frequency point (after FDR correction) inside the corresponding time-frequency window are highlighted in green.

**Figure 2.**
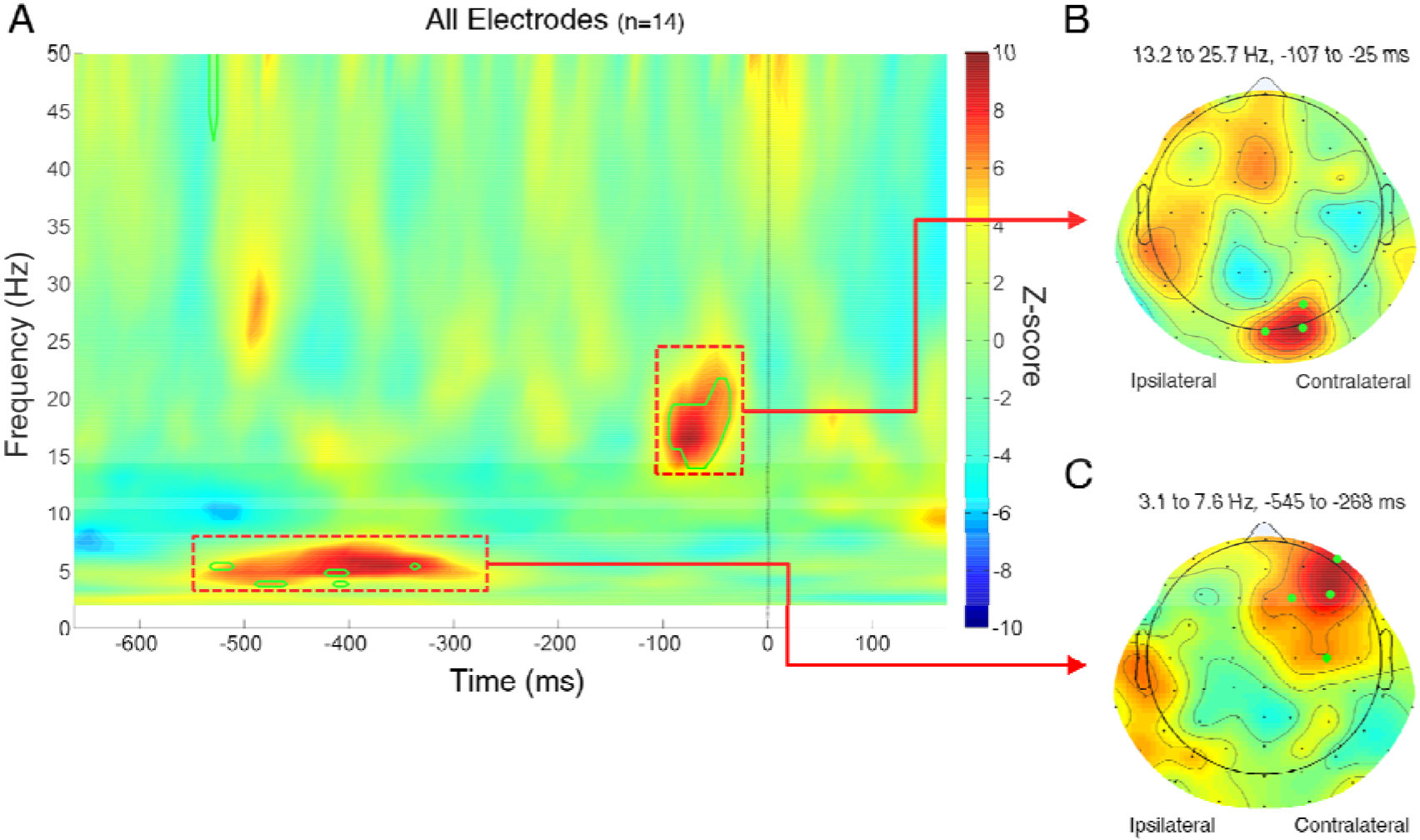
Pre-stimulus EEG phase predicts luminance judgment of 3D-shape disk vs. random-lines disk. (A) The relation between pre-stimulus phase and luminance judgment (our measure of predictive coding) is assessed using POP values (phase opposition product; see details in Methods). The time-frequency map is the z-score of observed POP values (combined across all subjects, datasets and electrodes), each value compared with a null-hypothesis distribution of 10,000 surrogate POP values (also combined across all subjects, datasets and electrodes) characterized by its mean and SD. Time 0 indicates stimulus onset. P-values were derived from a comparison of POP values against 80,000,000 surrogates, and corrected for multiple comparisons across all time points, frequencies and electrodes using the FDR method (FDR α = 0.05, corresponding to a P value threshold of 9.53 × 10^-6^). The green outlines mark the significant time-frequency regions (at least one significant electrode) after FDR correction. A significant relation is apparent between the effect of shape perception on luminance judgments and the EEG phase of ~5 Hz and ~16 Hz pre-stimulus oscillations. (B) Scalp topography of (z-scored) POP values around 16 Hz (frequency range 13.2 to 25.7 Hz; time range −107 to −25 ms). Electrodes highlighted in green have at least one significanttime-frequency point (after FDR correction) inside the corresponding red box. Due to our electrode-permutation procedure, in this topography the electrodes ipsilateral to the spatial choice are displayed on the left and those contralateral to the spatial choice are displayed on the right. Thus, the topography shows a contralateral occipital effect for the 16 Hz oscillation. (C) Same as B, but for the ~ 5 Hz oscillations (frequency range 3.1 to 7.6 Hz; time range −545 to −268 ms). The topography shows a contralateral frontal effect for the 5 Hz oscillation.

To quantify the influence of pre-stimulus oscillations on post-stimulus choice, we binned single trials according to the phase at the optimal time-frequency point (for the theta oscillation: −397 ms, 5.4Hz; for the beta oscillation: −68 ms, 16.5 Hz). Single trials were thus sorted in 13 phase bins based on the average phase of the significant electrodes for each oscillation (four frontal electrodes for the theta oscillation, three occipital electrodes for the beta oscillation). For each phase bin we then computed the post-stimulus choice probability of the 3D-shape disk. These choice probabilities were normalized by dividing them by the overall 3D-shape choice probability across all phase bins. For each experimental dataset (left-vs. right-side choice), phase bins were rotated such that the phase at which 3D-shape disk choice probability was largest was aligned to a phase angle of zero. As a result of this alignment, the 3D-shape choice probability is necessarily maximal at a phase angle of zero; therefore, the zero-phase bin was discarded from further analyses. For both frequencies, the 3D-shape disk choice probability monotonically decreased to a minimum at the opposite phase angle, confirming that pre-stimulus phase affected post-stimulus judgment (Figure 3). A one-way ANOVA showed that both pre-stimulus theta phase and pre-stimulus beta phase significantly modulated the 3D-shape disk choice probability (for theta oscillation, F_(11, 27)_ = 3.95, *p* = 2.23 × 10^-5^; for beta oscillation F_(11, 27)_=6.17, *p*=3.86 × 10^-9^). The magnitude of each effect was determined as the difference between the maximum and minimum 3D-shape disk choice probabilities across all phase bins. The frontal theta oscillation accounted for a difference of ~14% of the 3D-shape disk choice probability between phase bins, and the occipital beta oscillation accounted for a difference of ~19%.

**Figure 3.**
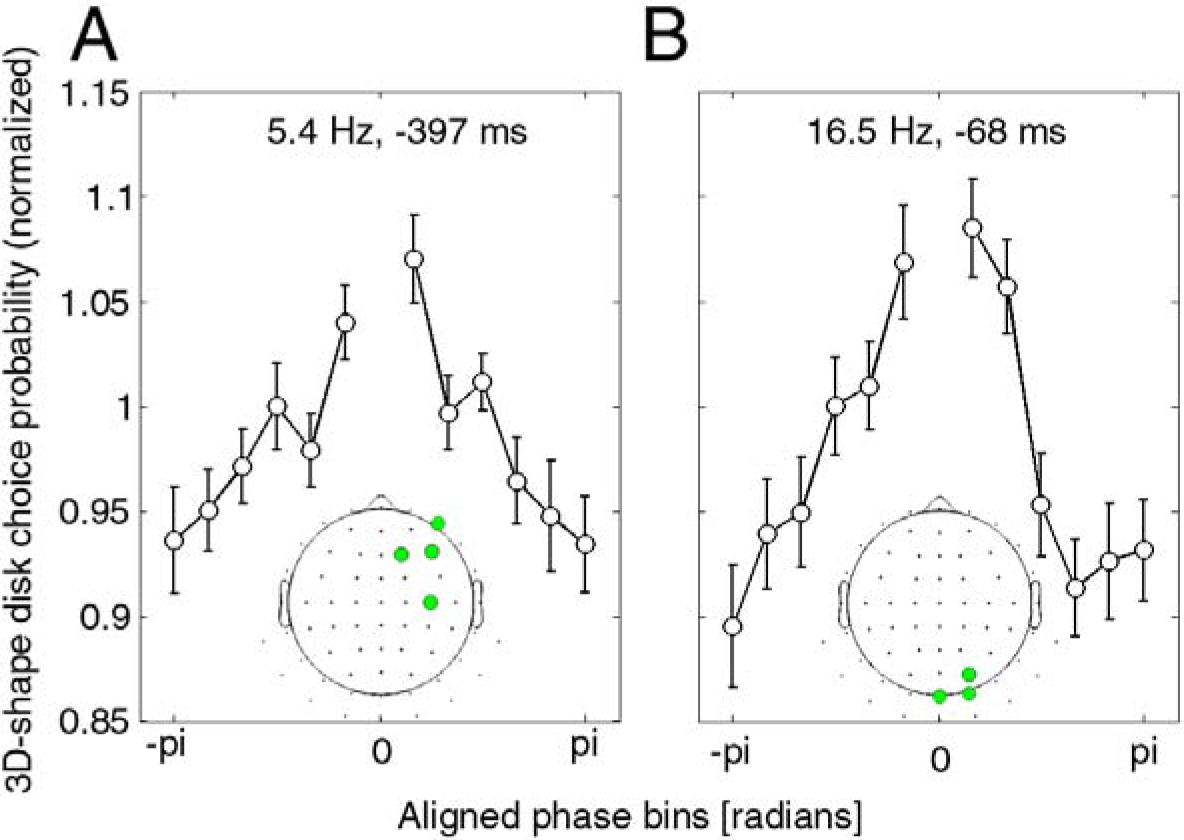
Normalized choice probability of 3D-shape disks as a function of pre-stimulus phase. (A) Relationship between frontal pre-stimulus theta phase and choice of 3D-shape disk as the brighter disk. Single trials were binned into 13 bins, centered on the maximal phase bin for each subject (central bin was then discarded). The curve indicates that the oscillatory phase of frontal electrodes (4 significant electrodes shown in inset topography) at 5.4 Hz and −397 ms modulates the luminance judgment by ~14%. Error bars represent SEM across subjects. (B) Same as A, but for the occipital beta pre-stimulus phase. The oscillatory phase of occipital electrodes (3 significant electrodes shown in inset topography) at 16.5 Hz and −68 ms modulates the luminance judgment by ~19%.

Because the time-frequency analysis relies on signal convolution with wavelet filters whose duration is non-negligible, one might wonder whether the observed pre-stimulus phase differences could actually be driven by stimulus-evoked activity. For example, at 16 Hz the above time-frequency analysis used a 250 ms time window (4 cycles, 125 ms from the past and 125 ms into the future); thus, significant phase effects observed at −67ms pre-stimulus may be contaminated by post-stimulus activity. To rule out such contamination, we repeated the POP time-frequency analysis with one-cycle wavelets at all frequencies, and compared the timing of pre-stimulus phase effects with the time-frequency region of possible poststimulus contamination, determined using the wavelet window length at each frequency (Figure 4). Both theta-and beta-frequency phase effects were replicated in this analysis, and were found to lie outside of the possible post-stimulus contamination zone.

**Figure 4.**
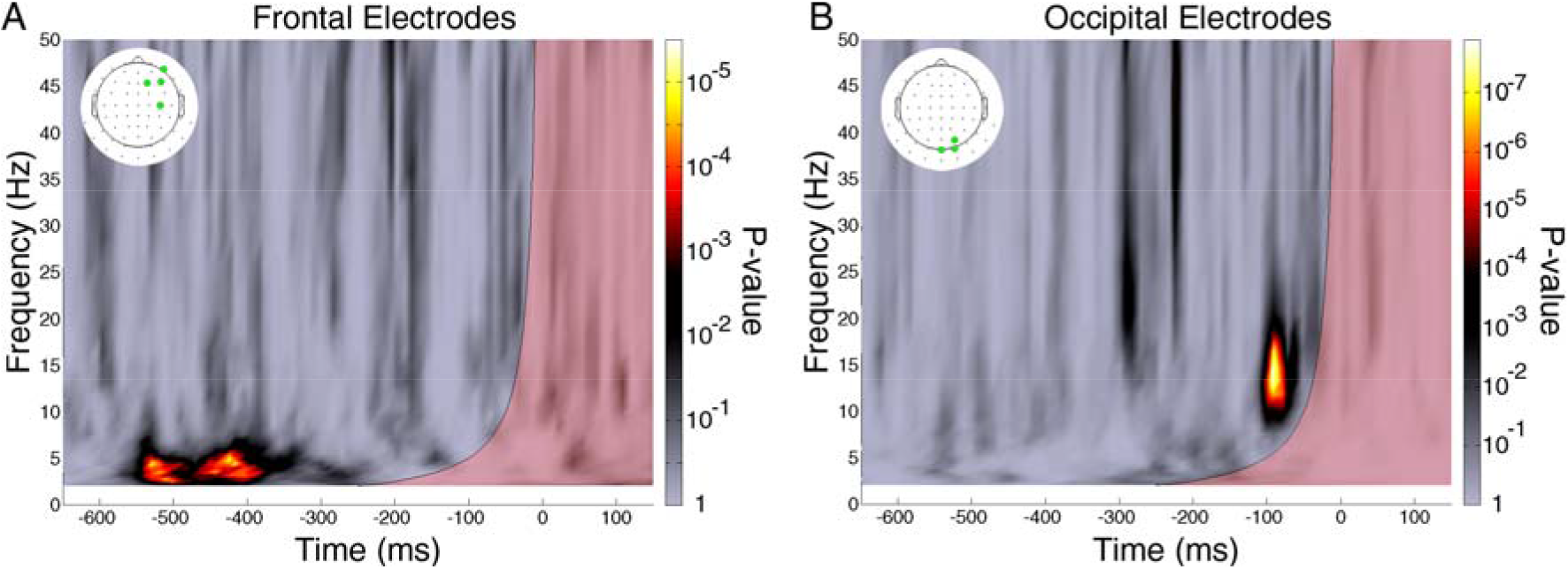
Significance of POP values in a one-cycle wavelet analysis. (A) P-value map of the POP values combined across previously identified frontal electrodes (green points in inset topography) for the theta-frequency phase effect. The P-values were calculated by comparing the observed POP values with 80,000,000 surrogates. The semi-transparent red area on the time-frequency map indicates the zone of possible contamination by post-stimulus activity (based on the wavelet window length at each frequency, centered on the time of stimulus onset, 0 ms). The previously observed theta-frequency phase effect lies outside of the contamination zone. (B) Same as A, but for the beta-frequency phase effect. The beta-frequency phase effect also lies outside the contamination zone. Altogether, these findings indicate that pre-stimulus phase differences are not caused by post-stimulus evoked activity.

We also ascertained that phase effects were not caused by any eye movement artifacts that may have survived our artifact rejection procedure. For example, the observed pre-stimulus phase differences could be thought to reflect different patterns of eye blink or saccades for different perceptual outcomes. Therefore, we applied our POP time-frequency analysis to the horizontal and vertical EOG signals. P-value maps (obtained by comparison of POP values against 80,000,000 surrogates) did not reveal any signs of systematic eye movements in either the theta-or the beta-frequency bands (Figure 5), ruling out an explanation of our prestimulus phase effects in terms of ocular artifacts.

**Figure 5.**
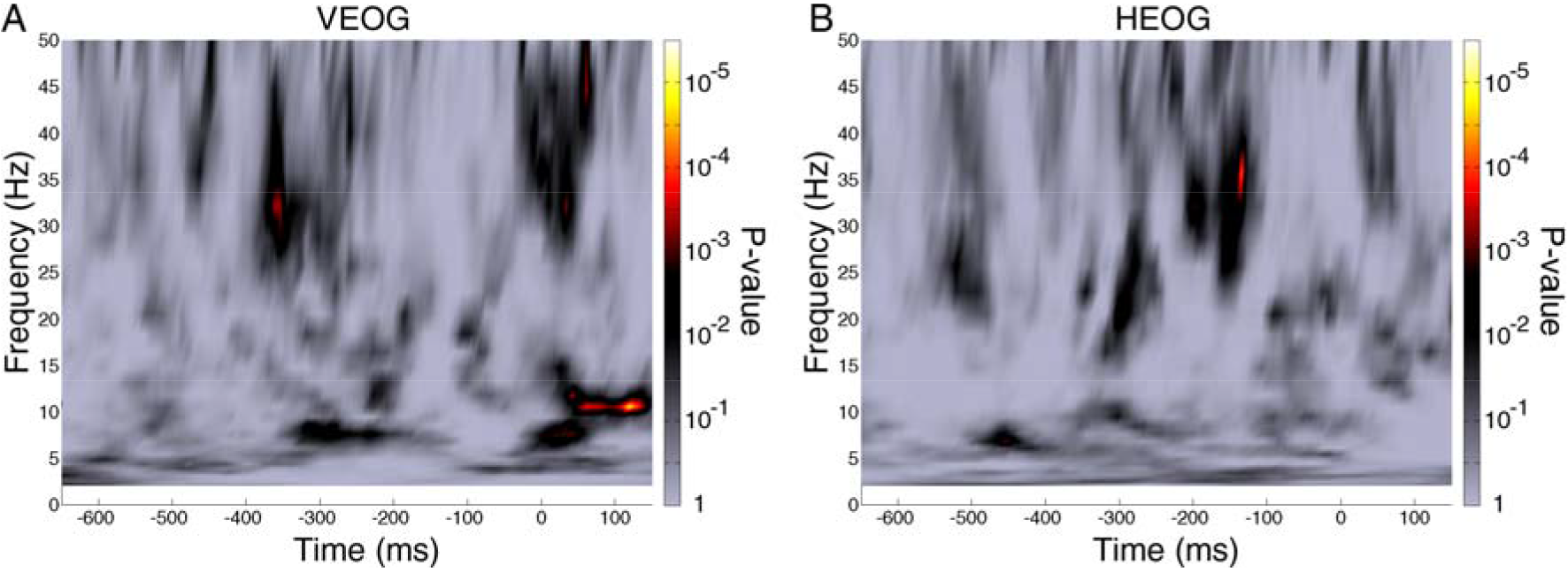
Significance of the POP values of VEOG and HEOG signals. (A) P-value map of the POP values for the VEOG. The P-values were calculated using a similar procedure as in the main analysis: comparing the observed POP values with 80,000,000 surrogate POP values. (B) Same as A, but for the HEOG. There was no significant pre-stimulus time-frequency window with significant POP values in either VEOG or HEOG, indicating that the observed phase effects were not due to ocular artifacts.

## Discussion

We investigated the temporal dynamics of predictive coding by exploring the relation between pre-stimulus oscillatory phase and the presumed trial-by-trial variations in poststimulus predictive feedback. We used 3D-shape outlines and random-lines versions of the same stimuli (as in one of the seminal predictive coding studies (Murray et al., 2002)) to induce different amounts of predictive feedback (Figure 1), and measured the corresponding effects on luminance judgment as trial-by-trial markers of the efficiency of predictive coding. (Feedforward processing of the 3D shape induced predictive feedback which increased the perceived brightness of the 3D-shape disk. The more efficient this predictive coding, the larger the effect of predictive feedback, and the brighter the 3D-shape disk Should be perceived.). Using a similar analysis method as in well-established studies of the relationship between pre-stimulus phase and post-stimulus behavior (Busch et al., 2009; VanRullen et al., 2011), we found that two pre-stimulus ongoing oscillations from different regions and frequencies could strongly influence the post-stimulus luminance judgment, and thus the post-stimulus predictive coding efficiency: a contralateral frontal theta oscillation and a contralateral occipital beta oscillation (Figure 2). The phase of the theta oscillation before stimulus onset could explain 14% of the luminance judgment difference while the phase of the beta oscillation could explain 19% (Figure 3). Control analyses ruled out contamination of the phase-behavior relationship by post-stimulus activity (Figure 4) or ocular artifacts (Figure 5). These results not only imply that predictive coding is a periodic process, but also reveal two periodicities with different sources. Since the occipital and frontal signals likely reflect activity from hierarchically lower and higher areas, respectively, and since predictive coding theory suggests that the brain sends back predictions from higher areas and sends prediction errors from lower areas, our results allow us to speculate on a possible temporal dynamic for predictive coding: predictions sent periodically at a theta frequency, prediction errors sent periodically at a beta frequency. Of course, more direct connectivity studies would be needed in the future to confirm these postulated roles for occipital and frontal signals.

The experimental paradigm used in this study takes advantage of the relationship between shape perception and predictive coding: 3D-shape outlines are assumed to generate more predictive feedback than the random-lines version of the same stimulus. Murray et al (2002) used similar stimuli to provide one of the first evidence of predictive coding: compared to the random-lines, 3D-shape outlines increased activity in the lateral occipital complex (LOC), but decreased it in primary visual cortex (V1), suggesting an increase of predictive feedback accompanied by a decrease in prediction errors (Clark, 2013; Murray et al., 2002). Here, we used the same paired stimuli as in the original study, placed them on two gray disks and asked subjects to judge the luminance of the disks. This luminance judgment, associated with perceived contrast, is likely to have a positive and monotonic relationship with neural activity in early visual cortex (Albrecht and Hamilton, 1982; Boynton et al., 1999; Dean, 1981; Goodyear and Menon, 1998). Thus, the variability in luminance judgment associated with the 3D-shape vs. random-lines stimuli could reflect trial-by-trial changes in the effect of predictive feedback on neural activity in early visual cortex.

Previous fMRI studies showed that shape perception could not only reduce (Murray et al., 2002), but also up-regulate neural activity in V1 (Kok and de Lange, 2014). Our own previous study found that 3D-shape disks were generally perceived brighter than random-lines disks, and that this effect could be attributed to predictive coding rather than attentional biases (Han and VanRullen, 2016, 2014). In the present study, we compensated for this net effect by adjusting the disks' luminance to obtain a 50% selection probability of 3D-shape/random-lines disks, and we focused on the remaining variability in luminance judgement as a trial-by-trial marker of the efficiency of predictive coding. On the other hand, systematic spatial biases (e.g. a general tendency to respond to the left or right stimulus) and/or trial-by-trial fluctuations in the direction of spatial attention can also be expected to affect the luminance judgement (Carrasco et al., 2004). As a matter of fact, spatial attention itself appears to involve a periodic process (Busch et al., 2009; Busch and VanRullen, 2010; Fiebelkorn et al., 2013; Landau and Fries, 2012) which could potentially influence the luminance judgement. We carefully avoided these potential confounding factors by dividing the trials into two datasets based on the post-stimulus spatial response (left/right) and performing the analysis within each dataset. If spatial attention biases, for example, were the only cause of the perceived luminance changes, the left-response dataset would pool all trials with a left-side attention bias (and similarly for the right-response dataset), and within each dataset pre-stimulus oscillatory phases would not bear any relation to post-stimulus luminance judgments. The existence of significant phase-behavior relationships in our analysis can therefore be safely attributed to predictive coding mechanisms rather than spatial attention or other biases.

Neurophysiological recordings have shown that feedforward and feedback may take advantage of oscillations in different frequencies. Laminar recordings showed that high-frequency oscillations are prominently generated in superficial layers and low-frequency oscillations in deep layers (Buffalo et al., 2011; Maier et al., 2010; Roopun et al., 2006). Since superficial and deep layers correspond respectively to the main sources of feedback and forward projections (Barbas and Rempel-Clower, 1997; Douglas and Martin, 2004; Wang, 2010), it follows that feedforward communication takes advantage of high-frequency oscillations and feedback takes advantage of low-frequency oscillations. A recent study with simultaneous recordings and micro-stimulation in different layers in V1 and V4 confirmed this notion (van Kerkoerle et al., 2014). Several authors have independently proposed that low-frequency oscillations send predictions via feedback, while high-frequency oscillations send prediction errors via feedforward (Arnal and Giraud, 2012; Bastos et al., 2012; Bauer et al., 2014; Fontolan et al., 2014; Todorovic et al., 2011; Yordanova et al., 2012). Our results provide support for this hypothesis at the EEG and behavioral level.

Our results also provide supportive evidence for the hypothesized functions of frontal theta-band and occipital beta-band oscillations. Our conclusions are in line with the notion that 5-10 Hz oscillations could contribute to “top-down” control (Jensen et al., 2012; VanRullen, 2013), which has already been suggested based on attentional phase effects on perception (Busch et al., 2009; Busch and VanRullen, 2010), reaction time (Drewes and VanRullen, 2011; Huang et al., 2015; Song et al., 2014) and perceptual variability in TMS-induced effects (Dugué et al., 2015, 2011). We found the origin of such theta periodicity in contralateral frontal electrodes, compatible with the involvement of frontal areas in the top-down controlling process (Summerfield and de Lange, 2014; Summerfield and Egner, 2009; Summerfield et al., 2006) and with the involvement of 5-10 Hz oscillations in this region (Phillips et al., 2014). On the other hand, local field potential (LFP) recordings showed that, in mammalian visual cortex, beta frequency oscillations are also prominent during the deployment of top-down control (Bekisz and Wróbel, 2003; Bosman et al., 2012; Buschman and Miller, 2007; Grothe et al., 2012; Lopes da Silva et al., 1970). Our findings of beta frequency phase effects on predictive feedback in the occipital area are concordant with such LFP results and suggest a valuable role for the beta frequency oscillations in predictive coding.

In summary, we measured the relation between pre-stimulus oscillations and a predictive feedback-induced effect to investigate the neural oscillations involved in predictive coding. We found that the pre-stimulus phases of frontal theta-frequency oscillations and occipital beta-frequency oscillations jointly determine post-stimulus subjective judgments. These results shed light on the temporal dynamics of predictive coding, and suggest a periodic predictive coding process with faster oscillations in lower areas and slower oscillations in higher areas.

## Acknowledgements

Biao Han is supported by China Scholarship Council; Rufin VanRullen is supported by an ERC Consolidator grant P-CYCLES number 614244.

## Competing interests

The authors declare no competing interests.

## Reference

Albrecht D.G., Hamilton D.B., 1982. Striate cortex of monkey and cat: contrast response function. J. Neurophysiol. 48, 217–237.

Alink A., Schwiedrzik C.M., Kohler A., Singer W., Muckli L., 2010. Stimulus Predictability Reduces Responses in Primary Visual Cortex. J Neurosci 30, 2960–2966.

Arnal L.H., Giraud A.L., 2012. Cortical oscillations and sensory predictions. Trends Cogn. Sci. 16, 390–398. doi:10.1016/j.tics.2012.05.003

Barbas H., Rempel-Clower, N., 1997. Cortical structure predicts the pattern of corticocortical connections. Cereb. Cortex 7, 635–646. doi:10.1093/cercor/7.7.635

Bastos A.M., Litvak V., Moran R., Bosman C., Fries P., Friston K.J., 2015. A DCM study of spectral asymmetries in feedforward and feedback connections between visual areas V1 and V4 in the monkey. Neuroimage 108, 460–475. doi:10.1016/j.neuroimage.2014.12.081

Bastos A.M., Usrey W.M., Adams R.A., Mangun G.R., Fries P., Friston K.J., 2012. Canonical microcircuits for predictive coding. Neuron 76, 695–711.

Bauer M., Stenner M.P., Friston K.J., Dolan R.J., 2014. Attentional Modulation of Alpha/Beta and Gamma Oscillations Reflect Functionally Distinct Processes. J Neurosci 34, 16117–16125.

Bekisz M., Wróbel, A., 2003. Attention-dependent coupling between beta activities recorded in the cat's thalamic and cortical representations of the central visual field. Eur. J. Neurosci. 17, 421–426. doi:10.1046/j.1460-9568.2003.02454.x

Bishop G., 1932. Cyclic changes in excitability of the optic pathway of the rabbit. Am. J. Physiol. Content.

Bosman, C. a., Schoffelen J.M., Brunet N., Oostenveld R., Bastos A.M., Womelsdorf T., Rubehn B., Stieglitz T., De Weerd, P., Fries P., 2012. Attentional Stimulus Selection through Selective Synchronization between Monkey Visual Areas. Neuron 75, 875–888. doi:10.1016/j.neuron.2012.06.037

Boynton G.M., Demb J.B., Glover G.H., Heeger D.J., 1999. Neuronal basis of contrast discrimination. Vision Res. 39, 257–269.

Buffalo, E. a, Fries P., Landman R., Buschman T.J., Desimone R., 2011. Laminar differences in gamma and alpha coherence in the ventral stream. Proc. Natl. Acad. Sci. U. S. A. 108, 11262–11267. doi:10.1073/pnas.1011284108

Busch, N. a, Dubois J., VanRullen R., 2009. The phase of ongoing EEG oscillations predicts visual perception. J. Neurosci. 29, 7869–7876. doi:10.1523/JNEUROSCI.0113-09.2009

Busch N.A., VanRullen R., 2010. Spontaneous EEG oscillations reveal periodic sampling of visual attention. Proc. Natl. Acad. Sci. U. S. A. 107, 16048–16053.

Buschman T.J., Miller E.K., 2007. Top-Down Versus Bottom-Up Control of Attention in the Prefrontal and Posterior Parietal Cortices. Science (80-.). 315, 1860–1862. doi:10.1126/science.1138071

Buzsáki, G., Draguhn A., 2004. Neuronal oscillations in cortical networks. Science 304, 1926–9. doi: 10.1126/science.1099745

Carrasco M.M., Carrasco M.M., Ling S., Ling S., Read S., Read S., 2004. Attention alters appearance. Nat. Neurosci. 7, 308–313.

Clark A., 2013. Whatever next? Predictive brains, situated agents, and the future of cognitive science. Behav. Brain Sci. 36, 181–204.

Dean A.F., 1981. The relationship between response amplitude and contrast for cat striate cortical neurones. J. Physiol. 318, 413–427.

Delorme A., Makeig S., 2004. EEGLAB: An open source toolbox for analysis of single-trial EEG dynamics including independent component analysis. J. Neurosci. Methods 134, 921. doi:10.1016/j.jneumeth.2003.10.009

Douglas R.J., Martin, K.A.C., 2004. Neuronal circuits of the neocortex. Annu. Rev. Neurosci. 27, 419–451.

Drewes J., VanRullen R., 2011. This Is the Rhythm of Your Eyes: The Phase of Ongoing Electroencephalogram Oscillations Modulates Saccadic Reaction Time. J Neurosci 31, 4698–4708.

Dugue L., Marque P., VanRullen R., 2015. Theta Oscillations Modulate Attentional Search Performance Periodically. J. Cogn. Neurosci. 945–958. doi:10.1162/jocn

Dugue L., Marque P., VanRullen R., 2011. The Phase of Ongoing Oscillations Mediates the Causal Relation between Brain Excitation and Visual Perception. J. Neurosci. 31, 11889–11893. doi:10.1523/JNEUROSCI.1161-11.2011

Egner T., Monti J.M., Summerfield C., 2010. Expectation and Surprise Determine Neural Population Responses in the Ventral Visual Stream. J Neurosci 30, 16601–16608.

Fiebelkorn I.C., Saalmann Y.B., Kastner S., 2013. Rhythmic sampling within and between objects despite sustained attention at a cued location. Curr. Biol. 23, 2553–8. doi:10.1016/j.cub.2013.10.063

Fontolan L., Morillon B., Liegeois-Chauvel, C., Giraud, A.-L., 2014. The contribution of frequency-specific activity to hierarchical information processing in the human auditory cortex. Nat. Commun. 5, 4694. doi:10.1038/ncomms5694

Fries P., Nikolic D., Singer W., 2007. The gamma cycle. Trends Neurosci. 30, 309–16. doi:10.1016/j.tins.2007.05.005

Friston K., 2005. A theory of cortical responses. Philos. Trans. R. Soc. Lond. B. Biol. Sci. 360, 815–836.

Goodyear B.G., Menon R.S., 1998. Effect of luminance contrast on BOLD fMRI response in human primary visual areas. J. Neurophysiol. 79, 2204–2207.

Grothe I., Neitzel S.D., Mandon S., Kreiter A.K., 2012. Switching neuronal inputs by differential modulations of gamma-band phase-coherence. J. Neurosci. 32, 16172–80. doi:10.1523/JNEUROSCI.0890-12.2012

Han B., VanRullen R., 2016. Shape perception enhances perceived contrast: evidence for excitatory predictive feedback? Sci. Rep. In-press.

Han B., VanRullen R., 2014. Predictive Coding of Shape Affects the Perceived Luminance of the Surrounding Region. J. Vis. 14, 72–72. doi:10.1167/14.10.72

Harrison L.M., Stephan K.E., Rees G., Friston K.J., 2007. Extra-classical receptive field effects measured in striate cortex with fMRI. Neuroimage 34, 1199–1208.

Huang Y., Chen L., Luo H., 2015. Behavioral Oscillation in Priming: Competing Perceptual Predictions Conveyed in Alternating Theta-Band Rhythms. J Neurosci 35, 2830–2837.

Jensen O., Bonnefond M., VanRullen R., 2012. An oscillatory mechanism for prioritizing salient unattended stimuli. Trends Cogn. Sci. 16, 200–206.

Kok P., de Lange F.P., 2014. Shape Perception Simultaneously Up-and Downregulates Neural Activity in the Primary Visual Cortex. Curr. Biol. 24, 1531–1535.

Lachaux J.P., Rodriguez E., Martinerie J., Varela F.J., 1999. Measuring phase synchrony in brain signals. Hum. Brain Mapp. 208, 194–208. doi:10.1002/(SICI)1097-0193(1999)8:4<194::AID-HBM4>3.0.CO;2-C

Landau A.N., Fries P., 2012. Attention samples stimuli rhythmically. Curr. Biol. 22, 1000–1004. doi:10.1016/j.cub.2012.03.054

Lopes da Silva, F.H., van Rotterdam, a, Storm van Leeuwen, W., Tielen, a M., 1970. Dynamic characteristics of visual evoked potentials in the dog. II. Beta frequency selectivity in evoked potentials and background activity. Electroencephalogr. Clin. Neurophysiol. 29, 260–268. doi:10.1016/0013-4694(70)90138-0

Maier A., Adams G.K., Aura C., Leopold, D. a, 2010. Distinct superficial and deep laminar domains of activity in the visual cortex during rest and stimulation. Front. Syst. Neurosci. 4, 1–11. doi:10.3389/fnsys.2010.00031

Murray S.O., Kersten D., Olshausen B.A., Schrater P., Woods D.L., 2002. Shape perception reduces activity in human primary visual cortex. Proc. Natl. Acad. Sci. U. S. A. 99,15164–15169.

Phillips J.M., Vinck M., Everling S., Womelsdorf T., 2014. A long-range fronto-parietal 5− to 10-Hz network predicts “top-down” controlled guidance in a task-switch paradigm. Cereb. Cortex 24, 1996–2008. doi:10.1093/cercor/bht050

Rao R.P., Ballard D.H., 1999. Predictive coding in the visual cortex: a functional interpretation of some extra-classical receptive-field effects. Nat. Neurosci. 2, 79–87.

Roopun A.K., Middleton S.J., Cunningham M.O., LeBeau F.E.N., Bibbig A., Whittington, M. a, Traub R.D., 2006. A beta2-frequency (20-30 Hz) oscillation in nonsynaptic networks of somatosensory cortex. Proc. Natl. Acad. Sci. U. S. A. 103, 15646–15650. doi:10.1073/pnas.0607443103

Song K., Meng M., Chen L., Zhou K., Luo H., 2014. Behavioral Oscillations in Attention: Rhythmic Alpha Pulses Mediated through Theta Band. J Neurosci 34, 4837–4844.

Summerfield C., de Lange, F.P., 2014. Expectation in perceptual decision making: neural and computational mechanisms. Nat. Rev. Neurosci.

Summerfield C., Egner T., 2009. Expectation (and attention) in visual cognition. Trends Cogn. Sci. 13, 403–409.

Summerfield C., Egner T., Greene M., Koechlin E., Mangels J., Hirsch J., 2006. Predictive codes for forthcoming perception in the frontal cortex. Science (80-.). 314, 1311–1314.

Summerfield C., Trittschuh E.H., Monti J.M., Mesulam, M.-M., Egner T., 2008. Neural repetition suppression reflects fulfilled perceptual expectations. Nat. Neurosci. 11, 1004–1006.

Todorovic A., van Ede, F., Maris E., de Lange, F.P., 2011. Prior Expectation Mediates Neural Adaptation to Repeated Sounds in the Auditory Cortex: An MEG Study. J. Neurosci. 31, 9118–9123.

van Kerkoerle T., Self M.W., Dagnino B., Gariel-Mathis, M.-A., Poort J., van der Togt, C., Roelfsema P.R., 2014. Alpha and gamma oscillations characterize feedback and feedforward processing in monkey visual cortex. Proc. Natl. Acad. Sci. U. S. A. 111, 14332–14341.

VanRullen R., 2013. Visual attention: a rhythmic process? Curr. Biol. 23, R1110–2.

VanRullen R., Busch N.A., Drewes J., Dubois J., 2011. Ongoing EEG Phase as a Trial-by-Trial Predictor of Perceptual and Attentional Variability. Front. Psychol. 2, 60. doi:10.3389/fpsyg.2011.00060

Wang, X.-J., 2010. Neurophysiological and computational principles of cortical rhythms in cognition. Physiol. Rev. 90, 1195–1268.

Yordanova J., Kolev V., Kirov R., 2012. Brain oscillations and predictive processing. Front. Psychol. 3, 1–2. doi:10.3389/fpsyg.2012.00416

